# Evaluation of a Computational Diagnostic for Epistasis in Plant Breeding Populations

**DOI:** 10.1101/044453

**Authors:** Reka Howard, William Beavis, Alicia Carriquiry

**Author notes:** Snedecor Hall, Ames IA 50011.

## Abstract

Previously we reported the inability of genomic prediction methods based on linear models to accurately predict trait values composed of an epistatic genetic architecture. We also reported non-parametric genomic prediction methods applied to the same data produced reasonably accurate predictions. The difference led us to propose analyses by paired parametric and non-parametric methods to the same data could be used as a diagnostic for epistatic genetic architectures in typical plant breeding populations. The suggested computational diagnostic was based on evaluation of 14 genomic prediction methods applied to eight sets of simulated conditions consisting of three factors, each with two levels. Because the potential set of factors that might affect accuracies of genomic predictions is unknown, there is a need for a systematic approach to identify combinations of factors that impact estimates of accuracy. Herein we propose the application of response surface methods to systematically identify conditions that maximize the difference between estimated accuracies of genomic prediction methods. The results indicate that genetic architecture and repeatability at their upper boundaries for complete epistasis and repeatability have the greatest influence on the differences between parametric and non-parametric estimated prediction accuracies. Further, the surface is very steep in the vicinity of the boundary conditions, indicating that the proposed diagnostic is of limited value for discovery of epistatic genetic architectures.

## INTRODUCTION

The application of molecular markers as surrogates for selection in genetic improvement of quantitative traits began over 35 years ago (Goodman and Stuber 1980; Goodman et al. 1980). With emergence of ubiquitous genomic marker technologies, Fernando and Grossman (1989) and Lande and Thompson (1990), proposed simple regression based marker assisted selection methods for genetic improvement of animals and plants. Anticipating the development of technologies capable of assaying tens of thousands of genomic markers on a few thousand phenotyped samples Meuwissen et al. (2001) proposed leveraging mixed models (Henderson 1975) to obtain predicted phenotypes for use in Genomic Selection (GS). Since their proposed methods a growing number of methods have been adopted to provide predicted phenotypes of individuals, or cultivars (Efron et al. 2004; Gianola et al. 2006; Maenhout et al. 2007; Park and Casella 2008; Piepho 2009; Usai et al. 2009; Pérez et al. 2010; Long et al. 2011). In the context of genetic gain, the impact of GS is to increase the selection intensity and decrease the time between cycles of selection with some reduction in the correlation between selection and response units. In aggregate the application of these methods are referred to as Genomic Selection (GS) or Genomic Prediction (GP) methods depending on whether the emphasis of the application is placed on selection, i.e., genetic gain or accuracy of prediction. Herein we refer to the methods as GP methods. Also, we distinguish GP methods as either parametric or non-parametric (Gianola et al. 2010), although the latter might be better described as machine learning methods.

With the proliferation of GP methods, there have also been published reviews (de los Campos et al. 2013; Gianola et al. 2010) and comparisons of the methods (VanRaden et al. 2009; Daetwyler et al. 2010; Jannink et al. 2010; de los Campos et al. 2010; Clark et al. 2011; Heslot et al. 2012 ; Riedelsheimer et al. 2012; Lorenz 2013; Howard et al. 2014; Thavamanikumar et al. 2015). The primary metrics for comparing GP methods include estimating Pearson’s correlation between predicted and actual phenotypes or estimated correlation divided by estimated heritability (Boddicker et al. 2014). Most of the comparative studies obtained estimated accuracies among GP methods using simulated data sets in which a true genotypic trait value is known.

Initial comparisons of GP methods focused on impacts of simulated numbers and magnitudes of additive quantitative trait loci (QTL) on the estimated accuracies of parametric methods. Most of these studies found that BayesB provided more accurate estimates of predicted values when the simulated numbers of QTLs are small but the magnitude of each QTL was simulated with large additive genetic effects (VanRaden et al. 2009; Daetwyler et al. 2010; Jannink et al. 2010). On the other hand, as the number of simulated QTLs increase and their individual magnitudes decrease all parametric methods (except linear regression) produce similar estimates of accuracy. Experimental studies on traits with a presumed known number of QTL tend to support the simulation studies (Clark et al. 2011; Thavamanikumar et al. 2015), although Riedelsheimer et al. (2012) did not find differences among parametric models in predicting traits with presumed large QTL effects. Machine learning techniques are also popular in GP to capture nonadditive effects. Heslot et al. (2012) compared several machine learning methods (reproducing kernel Hilbert space, neural network, random forest, and support vector machine) and techniques (bagging and boosting methods), but found no superior method in terms of accuracy of prediction.

To date parametric methods use models that ignore non-linear interactions, e.g., epistasis and genotype by environment interactions, in complex quantitative traits (de los Campos et al. 2010, Heffner et al. 2009, Hayes et al. 2009). These components are ignored primarily because the specific interactions are unknown and it is computationally difficult to search among all possible combinations of interactions (Moore 2009). Howard et al. (2014) compared estimates of accuracy among ten parametric and four non-parametric methods in a factorial design consisting of progeny types (F2 and backcross), heritability (0.3 and 0.7) and genetic architecture (additive and epistatic). From among the eight combinations of factors, only genetic architecture affected differences among estimated accuracies of the methods; for additive genetic architectures there were no significant differences of estimated accuracies among the methods, whereas non-parametric methods produced more accurate predictions than parametric methods for genetic architectures consisting of epistatic genetic effects. Indeed, parametric methods had no ability to predict the phenotypes, suggesting an analytical diagnostic could reveal underlying unknown genetic architecture in experimental data.

In addition to genetic architecture of the trait, there are other factors that may affect differences of estimated accuracies. These include resource allocation in terms of sample size, number of families, number of lines per family, numbers of marker loci, number of replicated field trials (Lorenz 2013), population structure in terms of numbers and relatedness among families (Hickey et al. 2015), proportion of phenotypic variability that can be attributed to additive and non-linear components, etc. To date, some of these additional factors have been investigated, but in an ad hoc manner. A more systematic approach is needed to find the conditions under which various GP methods will produce the best predictions.

Response Surface Methods (RSM) were developed and introduced by Box and Wilson (1951) to experimentally search for combinations of factors that will maximize a response metric. The goal is to reduce the number of experimental treatment combinations needed to find the maximum desirable response (Naylor 1969). Myers et al. (1989) summarized the extensive applications of RSM to engineering systems. Bezerra et al. (2008) summarized applications in analytical chemistry and more recently RSM have been applied to pharmaceutical production (Koyamada et al. 2004), water purification (Nair et al. 2014), food processing (Afoakwa et al. 2014) and fermentation systems (Zhang et al. 2010). Importantly for our consideration are applications of RSM to development of machine learning methods (Gonen and Alpaydin 2011). Despite clear efficiencies of RSM, we have found no evidence of their application in development and evaluation of GP methods. Herein we introduce RSM as an effective and efficient means to find situations in which application of both parametric and non-parametric GP methods can be used to diagnose the underlying genetic architecture of a trait. Specifically, in response to a proposed computational diagnostics (Howard et al. 2014) our goal is to find the combination of factors and factor levels for which the nonparametric methods outperform the parametric methods in terms of prediction accuracy.

## MATERIALS METHODS

### Response Metric and Genomic Prediction Methods

We define prediction accuracy as the correlation between the predicted and the true simulated phenotypes. Our primary response of interest is the difference between prediction accuracies generated by parametric and non-parametric methods. BLUP describes the best linear unbiased properties used to estimate fixed effects and predict random effects in a mixed linear model (Henderson et al. 1959, Henderson 1963). In the context of GP it consists of a random effect parameter for the realized relationship matrix that is estimated with marker genotypic scores (Bernardo 1994; Hayes et al. 2009). Because Howard et al. (2014) found that BLUP values were as accurate as other parametric GP methods for backcross and F2 populations and BLUP has been implemented in the R package rrBLUP (Endelman 2011) we chose it to represent the parametric GP methods. The Support Vector Machine (SVM) is a nonparametric machine learning technique that can model the relationship between the marker values and the phenotypes using a linear or a nonlinear function (Vapnik 1995; Hastie et al. 2009; Howard et al. 2014). Because Howard et al. (2014) found that predicted values from the SVM were as accurate as other non-parametric GP methods for backcross and F2 populations and it has been implemented in the R package kernlab (Karatzoglou et al. 2004) we chose it to represent the non-parametric GP methods.

### Simulation Design

We used cross validation, where we divided simulated data into training sets and testing sets. The training sets were used to fit the models, and the testing sets were used to calculate the accuracy of prediction for each GP method. We determined the differential between prediction accuracies for each GP prediction method as a function of the number of QTL, interactions among QTL, heritability, sample size, and number of markers. Because previous analyses (Howard et al. 2014) found little differential among prediction accuracies for GP methods associated with differences among types of plant breeding populations (F2, BC, DH and RIL) we simulated only BC families. Simulations were conducted using the R package QTL Bayesian Interval Mapping (Yandell et al. 2012). R can be accessed at http://www.r-project.org, the qtlbim package can be obtained by library qtlbim (Yandell et al. 2007) and the reference manual at http://cran.r-project.org/web/packages/qtlbim/qtlbim.pdf. The simulated genome had 10 chromosomes, each having the same length. The markers were distributed throughout the genome in such a way that each chromosome had the same number of markers equally spaced along the length of each chromosome. Likewise, QTL were distributed uniformly among the chromosomes. Cooper et al. (2002) discussed the potential contribution of multi-way gene interactions to an adaptive surface: if the average number of genetic interactions is greater than two the adaptive surface is becomes sufficiently rough that responses to selection are limited. Because plant breeding populations have responded to selection for at least 70 years, we simulated only two-way gene interactions among QTL when epistasis was included in the simulation model. No missing genotypic values and no missing phenotypic values were simulated in any data sets. Errors associated with phenotypic values were sampled from a normal distribution. For each combination of factors (Table 1) 20 replicates were simulated. Each replicate was subsequently divided into 25 pairs of training and testing data sets by sampling without replacement 80% of the simulated BC family members to provide a training set and used the remaining 20% of the members to provide a testing set. In total, this resulted in 500 replicates for each combination of factors.

**◼ Table 1.** Specification of the two levels of five factors include *n*, number of segregating progeny, *m*, marker number, QTL number, proportion of genetic variance due to epistasis and heritability. Epistasis 0 means that all of the genetic variance is additive variance, 0.5 epistasis means that half of the genetic variance is additive and the other half is epistatic.

| Factor | Level 1 | Level 2 |
| --- | --- | --- |
| n | 200 | 1000 |
| m | 50 | 400 |
| qtl | 10 | 50 |
| epi | 0 | 0.5 |
| h | 0.2 | 0.5 |

### RSM Strategy

To estimate prediction accuracies for all factor combinations consisting of two levels per factor would require a minimum of 2^*p*^ treatment combinations in a full factorial treatment design. When p is large and the range of possible values of each factor also is large, finding the combination of the *p* factors that define the response surface increases dramatically. For example if five levels per factor were needed to identify multiple peaks on the surface, then a minimum of 5^*p*^ factor combinations would be needed. In addition if statistical inference about the variability in the estimated response is desired, then at least some of the treatment combinations need to be replicated. Fortunately, our goal is not to describe the response surface, rather to maximize the differential response between types of GP methods. Thus, there is no need to investigate all possible sets of factor combinations, rather to find the regions where the differential between types of GP methods is largest.

Let *ph*_*i*_ denote the true phenotypic values at factor combination *i* and let *pĥ*_*i*_ denote the estimated phenotypic values at factor combination *i*. The response, *y* depends on a set of design variables *x*_*ind*_, *x*_*m*_, *x*_*QTL*_, *x*_*epi*_, and *x*_*h*_, where *x*_*ind*_ is the number of individuals in the simulated backcross population, *x*_*m*_ is the number of markers, *x*_*QTL*_ is the number of QTL, *x*_*epi*_ is the proportion of genotypic variability due to interactions among QTL (epistasis), and *x*_*h*_ is the proportion of phenotypic variability due to genotypic variability (heritability). The model can be written as

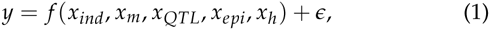

where *y* is the response, *ƒ* is the unknown, possibly complex response function, which depends on the design variables *x*_*ind*_, *x*_*m*_, *x*_*QTL*_, *x*_*epi*_, and *x*_*h*_, and *∊* ~ iid *N*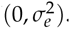. The expected value of the response function can be written as

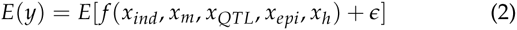

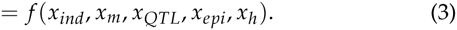

Because there are five factors under consideration, where each factor could have multiple, *m*, levels possible for each of these continuous parameters, a full factorial design would require *m*^5^ experimental treatment combinations. To search the space defined by these parameters for the maximum differential among GP methods we adopted a steepest ascent strategy (Myers, 1976). In the steepest ascent approach an initial experiment is designed such that only two of the *m* possible values for each factor is sampled and many of the potential interactions among factors are confounded. Specifically our initial experiment was a half fractional factorial design consisting of 2^*p*-1^ = 16 factor combinations, where each combination was represented by 500 replicates to estimate the differential response between GP methods. The initial values for each factor combination are listed in Table 1.

Initially, we use a first-order polynomial to approximate the response function, *ƒ*, so that

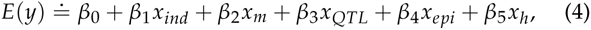

where *β*_0_ is the intercept, *β*_1_, is the regression coefficient associated with the number of individuals, *β*_2_ is the regression coefficient associated with the number of markers, *β*_3_ is the regression coefficient associated with the number of QTL, *β*_4_ is the regression coefficient associated with the proportion of genetic variation due to epistasis, and *β*_5_ is the regression coefficient associated with heritability. After completing the analyses of the initial fractional factorial the next set of conditions for a second set of conditions is determined by calculating a ‘base value’ from which increments to each factor that maximize change in response can be calculated. The base value is calculated as the average of the estimated low and the high levels of the factors. The increment is calculated as the product of 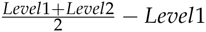 and the corresponding movement values (Myers et al. 1989) which determine which direction and magnitude we have to move on the response surface for each factor. This process is then repeated until estimated increments are less than some arbitrarily small value |*i*_*r*_| for experimental condition r.

### Data Availability

The simulated data, the R code used for simulating the data, and the R code for the parametric BLUP and nonparametric SVM can be found at http://gfspopgen.agron.iastate.edu/Supplemen-tary%20Resources1.html.

## RESULTS

Averages of 500 estimated prediction accuracies for BLUP, SVM and the difference between the methods for the initial 16 combinations (Table 2) were analyzed using analysis of variance. The results indicated that the best fit model included no interactions among factors with the following estimates of the differences between low and high values for each of the initial factors:

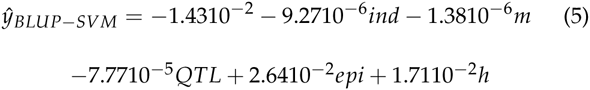

**◼ Table 2.** Mean accuracy of BLUP, mean accuracy of SVM, and the response (difference of mean accuracy of SVM and mean accuracy of BLUP) for 16 treatment combinations.

| Treatment Combination | BLUP | SVM | Diff |
| --- | --- | --- | --- |
| 200 ind, 50 m, 10 QTL, 0 epi, 0.5 h | 0.62 | 0.59 | -0.02 |
| 1000 ind, 50 m, 10 QTL, 0 epi, 0.2 h | 0.40 | 0.37 | -0.03 |
| 200 ind, 400 m, 10 QTL, 0 epi, 0.2 h | 0.23 | 0.22 | -0.01 |
| 1000 ind, 400 m, 10 QTL, 0 epi, 0.5 h | 0.63 | 0.61 | -0.02 |
| 200 ind, 50 m, 50 QTL, 0 epi, 0.2 h | 0.05 | 0.05 | 0.00 |
| 1000 ind, 50 m, 50 QTL, 0 epi, 0.5 h | 0.29 | 0.27 | -0.03 |
| 200 ind, 400 m, 50 QTL, 0 epi, 0.5 h | 0.28 | 0.28 | -0.01 |
| 1000 ind, 400 m, 50 QTL, 0 epi, 0.2 h | 0.23 | 0.21 | -0.02 |
| 200 ind, 50 m, 10 QTL, 0.5 epi, 0.2 h | 0.14 | 0.13 | -0.01 |
| 1000 ind, 50 m, 10 QTL, 0.5 epi, 0.5 h | 0.48 | 0.51 | 0.03 |
| 200 ind, 400 m, 10 QTL, 0.5 epi, 0.5 h | 0.27 | 0.28 | 0.01 |
| 1000 ind, 400 m, 10 QTL, 0.5 epi, 0.2 h | 0.22 | 0.19 | -0.03 |
| 200 ind, 50 m, 50 QTL, 0.5 epi, 0.5 h | 0.06 | 0.05 | -0.01 |
| 1000 ind, 50 m, 50 QTL, 0.5 epi, 0.2 h | 0.08 | 0.07 | -0.01 |
| 200 ind, 400 m, 50 QTL, 0.5 epi, 0.2 h | 0.05 | 0.05 | 0.00 |
| 1000 ind, 400 m, 50 QTL, 0.5 epi, 0.5 h | 0.25 | 0.23 | -0.02 |

Note that the model had the largest estimated coefficient for epistasis, thus changes to this factor will have the greatest impact on increasing estimated differences between predictive accuracies. We chose a basis to be 1% of epistasis corresponding to 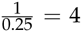 design units. The basis influenced the increment (Δ) of the other factors, and the change values of the four other factors are:

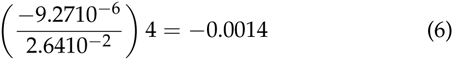

for individuals,

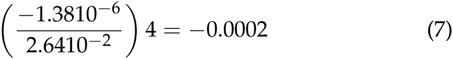

for markers,

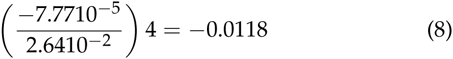

for QTL,

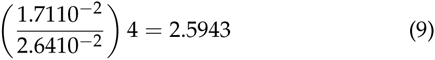

for heritability. The resulting increment for the number of individuals, number of QTL, and the number of markers were close to zero, which indicated that the levels for these variables did not need to be changed for the next set of experimental runs. The proportion of genetic variability due to epistasis and magnitude of heritability are the only factors that needed to change in order to expect significant changes on the response surface. A Base+3Δ will produce values that are near the boundary limits for epistasis, and heritability. It is not expected that further changes to *n*, the number of individuals, *m*, the number of markers, and the number of QTL will improve the differential *y* between estimated prediction accuracies for types of GP methods.

The response values from additional experiments are larger, in most cases, than the response values were for the 16 initial treatment combinations (table 3). The maximum difference in estimated prediction accuracies of GP methods was found to consist of 597 backcrossed progeny, 225, markers, and a genetic architecture consisting of 29 QTL in which all of the genetic variability is due to epistasis and the genetic variability is responsible for 100% of the phenotypic variability.

**◼ Table 3.**
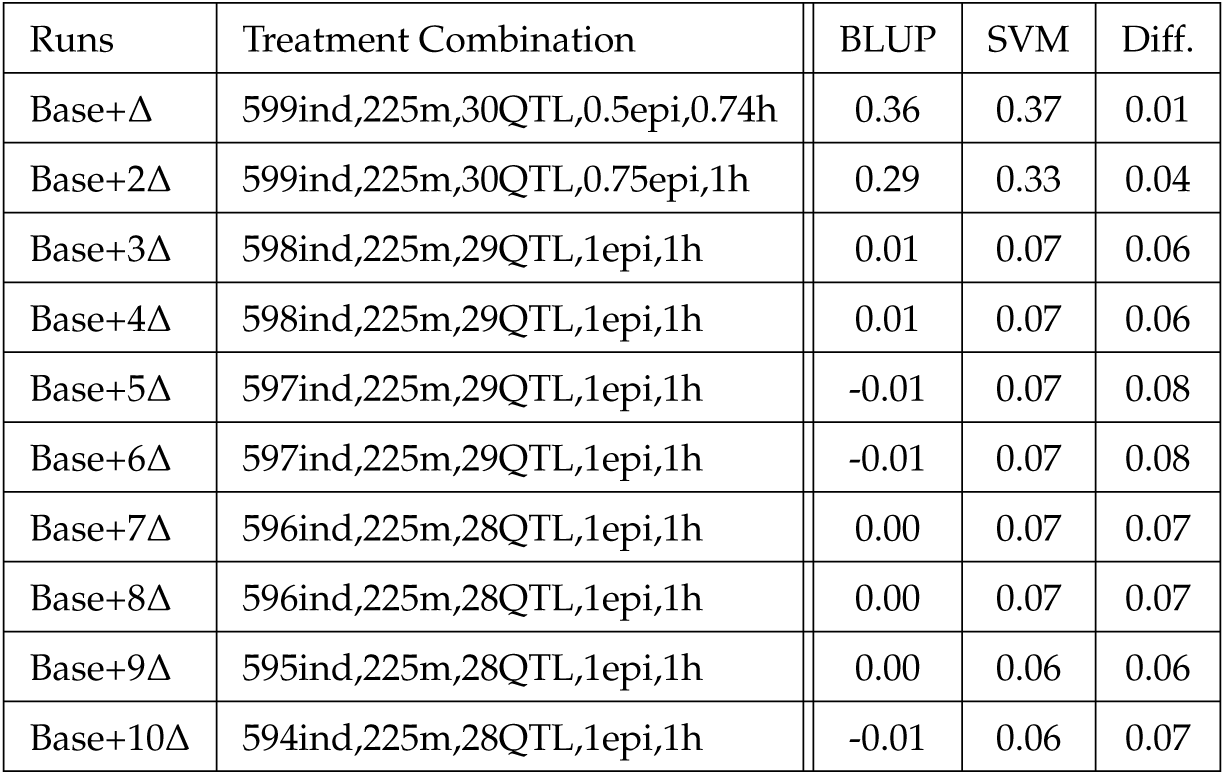
Mean accuracy of BLUP, mean accuracy of SVM, and the response (difference of mean accuracy of SVM and mean accuracy of BLUP) for the additional treatment combinations for the additional runs.

## DISCUSSION

Howard et al. (2014) proposed a computational diagnostic to reveal epistatic architectures based on estimated differential prediction accuracies generated by parametric and nonparametric GP methods. Their study reaffirmed that application of methods using incorrect linear models produce results that are worse than machine learning methods, while methods that do not require model specification perform almost as well as methods with correctly specified linear models. The challenge facing plant breeders is that the underlying genetic architecture of complex and quantitative traits are unknown. To our knowledge only one experimental study has supported the proposed computational diagnostic of epistasis. Clowers et al. (2010) found that parametric genomic prediction methods were not predictive of response to cold shock in Drosophila, whereas nonparametric methods produced reasonably accurate predictions. Since the underlying genetic architecture for this highly heritable trait is known to consist of more than a dozen interacting genetic loci, their application of a computational diagnostic to data representing a reasonable number of markers and sampled progeny were able to reveal an epistatic genetic architecture for the trait.

The goal of the research reported herein was to identify conditions under which the proposed computational diagnostic could be applied and interpreted. The critical first step was to establish objective measurable criteria that matched the goal. Specifically, we investigated combinations of sample size, number of markers, number of QTL, proportion of genetic variability due to interactions among QTL and proportion of phenotypic variability due to genotypic variability (heritability) for which nonparametric GP methods outperform parametric methods. An alternative objective might have been to identify the combination of factors that would maximize accuracy of predictions for each type of GP method, but such an alternative would not have addressed the goal. Because each of the five variable factors could assume an infinite number of possible positive values a systematic approach was needed for the investigation. To minimize the number of simulated experimental combinations, we chose to use the steepest ascent strategy from among Response Surface Methods.

The results indicate that the initial upper values for number of progeny, number of markers and number of QTL did not need much change in order to maximize the response. Fernando and Garrick have learned that if the training population is small (<1,000) or large (>100,000) GP methods produce asymptotically equivalent prediction accuracies, but sample sizes between these extremes will result in differential prediction accuracies (personal communication). Because the reproductive biology of most crop species is not capable of producing samples consisting of tens of thousands of (BC, DH or RIL) progeny such sample sizes would not be considered unless data from multiple families are combined (Hickey et al., 2014). We hypothesize that variability of relationships among families will affect differential responses among GP methods for these intermediate numbers of progeny in the training data sets.

Heritability and the proportion of genetic architecture due to epistasis with values at the upper boundary of 1.0 have the greatest influence on the differences between parametric and nonparametric estimated prediction accuracies. Note that estimated prediction accuracies reported herein are not identical to results previously reported (Howard et al. 2014). This is likely due to differences in simulated genomic structures between the studies. Howard et al., (2014) simulated variable length chromosomes, and herein we simulated uniform length chromosomes. Consequently, the relative genomic locations and linkage disequilibria among the simulated QTL were not the same in both studies. Future work should examine how genomic structure and/or linkage disequilibria can influence differential estimated prediction accuracies of GP methods.

We also learned that the surface is very steep in the vicinity of the boundary conditions, (data not shown) indicating that the proposed diagnostic is probably of limited value for discovery of epistatic genetic architectures for most traits of agronomic importance. Virtually all molecular genetic models of genetic expression include regulatory motifs that depend on expression of other genetic loci in cell signaling networks (Eungdamrong and Iyengar 2004), while estimates of genetic variance components for most agronomic traits are ascribed to additive genetic effects. The seeming paradox between molecular and quantitative genetic models has been the subject of considerable modeling efforts (Cheverud and Routman 1995; Aylor and Zeng 2008) that have yet to resolve the issue. And, results reported herein indicate that the proposed computational diagnostic by Howard et al. (2014) will not contribute to resolution of the issue except in extreme situations where contributions of epistasis and heritability are at their respective boundary conditions.

We did not investigate a comprehensive set of factors that could affect differential prediction accuracies among GP methods. For example, we did not include variability among factors such as genetic relationships among multiple families (Hickey et al. 2014), disequilibrium among QTLs and genotype by environment interaction effects used in training and testing populations. Thus, we found a local maximum for the response and future work will be needed to determine if this is a global maximum. The next set of evaluations should include factors such as family structure (eg. multiple families versus single family), multiple environments, family relationship between individuals in the training and testing sets, disequilibrium among QTL, and chromosomal distribution of the QTL. We hypothesize that the difference between parametric linear models and non-parametric machine learning methods in terms of prediction accuracy will grow larger on a more complex surface. However, the interpretation of the diagnostic will not be based solely on genetic architecture such as we found in our local optima, rather it will be confounded with a number of possible non-linear and probably interacting factors.

We demonstrated that a steepest ascent RSM can be used to find the conditions under which the differential is maximized. Potential hundreds of combinations of non-linear interacting factors reduced to investigation of only *n* experimental combinations. Even though we illustrated the implementation of RSM using a GP example, the methodology can be applied to development of any data analysis technique. We hypothesize that application of RSM to development of analytics for biological ‘big data’ challenges from genomes to field systems will be effective and efficient at finding the optimal conditions for employing the methods. Indeed Gonen and Alpaydin (2011) have demonstrated the use of RSM in development of a machine learning method.

